# A metabolomics study of ascorbic acid-induced *in situ* freezing tolerance in spinach (*Spinacia oleracea* L.)

**DOI:** 10.1101/2020.01.23.916973

**Authors:** Kyungwon Min, Keting Chen, Rajeev Arora

## Abstract

Freeze-thaw stress is one of the major environmental constraints that limit plant growth and reduces productivity and quality. Plants exhibit a variety of cellular dysfunction following freeze-thaw stress, including accumulation of reactive oxygen species (ROS). This means that enhancement of antioxidant capacity by exogenous application of antioxidants could potentially be one of major strategies for improving freezing tolerance (FT) of plants. Exogenous application of ascorbic acid (AsA), as an antioxidant, has been shown to improve plant tolerance against abiotic stresses but its effect on FT has not been investigated. We evaluated the effect of AsA-feeding on FT of spinach (*Spinacia oleracea* L.) at whole-plant and excised leaf level, and conducted metabolite profiling of leaves before and after AsA-treatment to explore metabolic explanation for change in FT. AsA-application did not impede leaf-growth; instead slightly promoted it. Temperature-controlled freeze-thaw tests revealed AsA-fed plants were more freezing tolerant as indicated by: 1) less visual damage/mortality; 2) lower ion-leakage; and 3) less oxidative-injury, lower abundance of free radicals (O_2_ ^•−^ and H_2_O_2_). Comparative leaf metabolic profiling revealed clear separation of metabolic phenotypes for control *vs*. AsA-fed leaves. Specifically, AsA-fed leaves had greater abundance of antioxidants (AsA, glutathione, alpha-& gamma-tocopherol) and compatible solutes (proline, galactinol, myo-inositol). AsA-fed leaves also had higher activity of antioxidant enzymes (superoxide dismutase, ascorbate peroxidase, catalase). These changes, together, may improve FT via alleviating freeze-induced oxidative stress as well as protecting membranes from freeze-desiccation. Additionally, improved FT by AsA-feeding may potentially include enhanced cell wall/lignin augmentation and bolstered secondary metabolism as indicated by diminished level of phenylalanine and increased abundance of branched amino acids, respectively.

## 1 INTRODUCTION

Sub-freezing temperatures are the major environmental constraint affecting crop performance and limiting plant distribution. This provides ample incentive to improve plants’ freezing tolerance (FT). Freeze thaw-injured tissues undergo various cellular dysfunctions. Thus far, two of the most studied loci of such injury are 1) leakage of cellular solutes, i.e. physico-molecular perturbations in cell membranes, and 2) oxidative injury to macromolecules due to cellular accumulation of reactive oxygen species (ROS; e.g. superoxide, singlet oxygen etc.) (Kendall & McKersie, 1989; Mittler, 2002; Min, Chen, & Arora, 2014; Arora, 2018). Hence, detoxification of excess ROS is believed to be one of the major strategies of frost survival (McKersie et al., 1993; McKersie, Bowley, & Jones, 1999).

Certain plants from temperate region have an ability to increase their FT, via a process called cold acclimation, when exposed to cold temperature (Thomashow, 2010). This involves a myriad of adjustments at physiological, biochemical, and metabolic levels, including an upregulation or accumulation of enzymatic and/or non-enzymatic antioxidants (Xin & Browse, 2000; Thomashow, 2010). This suggests enhancement of antioxidant capacity by exogenous application of antioxidants could potentially be an intervention strategy to increase plants’ FT. Ascorbic acid (AsA) is a well-known water-soluble antioxidant involved in ascorbate-glutathione cycle, especially as a substrate for ascorbate peroxidase (APX) which is responsible for converting H_2_O_2_ into H_2_O (Smirnoff, 2000; Foyer & Noctor, 2011). Research has shown exogenous AsA to improve plant tolerance against salt, drought, and chilling (Amin, Mahleghah, Mahmood, & Hossein, 2009; Azzedine, Gherroucha, & Baka, 2011; Ahmad, Basra, & Wahid, 2014; Akram, Shafiq, & Ashraf, 2017). But no study, to our knowledge, exists on the effect of AsA on FT of whole plants. Moreover, a comprehensive study of metabolome changes induced by AsA feeding of tissues could provide additional insight into biochemical mechanism and *in vivo* role of AsA-induced stress tolerance, including FT. These studies may also lead to identification of beneficial metabolites *vis-à-vis* FT enhancement.

In the present study, our main goals are twofold to: (1) investigate the effect of AsA-feeding on FT of spinach seedlings at the whole-plant as well as excised-leaf level, and (2) explore metabolome changes induced by AsA-treatment using gas chromatography-mass spectrometry (GC-MS). We used spinach as a model because of its moderate constitutive FT allowing sufficient range of freezing treatment temperatures for the present study and our previous experience with this system (Chen & Arora, 2014; Shin, Min, & Arora, 2018; Min, Showman, Perera, & Arora, 2018).Visual estimation and ion-leakage test were used to evaluate AsA-induced FT following *in situ* freezing test. Other physiological parameters, i.e. histochemical detection of ROS, activity of antioxidant enzymes, and leaf content of glutathione (GSH) were also determined for untreated control and AsA-fed tissues.

## 2 MATERIALS AND METHODS

### 2.1 Plant materials

Spinach seedlings were grown as described previously (Min, Showman, Perera, & Arora, 2018). Briefly, seeds of ‘Reflect’, a F_1_ hybrid cultivar (Johnny’s selected seeds, Inc., Winslow, ME, USA) were sown in plug flats filled with Sunshine LC-1 mix (Seba Beach, Alberta, Canada) and placed in growth chambers at 15/15 °C (D/N) with 12-h photoperiod under average PAR of ∼300 µmol m^-2^ s^-1^ at plant height provided by incandescent and fluorescent lamps. Seedlings were watered as needed via sub-irrigation (approximately, 5-d interval). After two weeks from the sowing, temperature in chambers was elevated to 20/18 °C (D/N), and seedlings were sub-fertigated only once with either 300 ppm EXCEL (Everris NA Inc., Dublin, OH, USA) nutrient solution (hereafter referred to as F-control) or with 0.5 and 1.0 mM AsA treatment made with 300 ppm EXCEL as solvent. About 24-day-old spinach seedlings, i.e. 10-d after fertigation treatments, were used for experiments as described below.

### 2.2 Growth measurement

Leaf growth was evaluated by measuring fresh weight (FW), dry weight (DW), and leaf area of F-control and AsA-fed leaves. Briefly, 7 to12 pairs of leaves (total 14 to 24 leaves) per treatment were first used to measure leaf area using LI-3100 Area Meter (LI-COR, Inc., Lincoln, NE, USA), quickly followed by the measurement of FW on the same leaves. DW was measured following oven-drying leaves at 75 ± 1 °C for 72-h. Water content was calculated on FW basis. Data of leaf-growth across five biological replications (14 to 24 leaves per biological replicate) were pooled to calculate the representative treatment means with standard errors. Mean differences between treatments were analyzed by least significant difference (LSD) test.

### 2.3 Freezing tolerance measurement

#### 2.3.1 *In situ* freezing test

Temperature-controlled, whole-plant freezing protocol was used, as described by Min, Showman, Perera, & Arora (2018), to compere FT between F-control and AsA-fed seedlings. Three plug flats – one of F-control and the other two with 0.5 or 1.0 mM AsA-fed plants - were transferred to a freezing chamber (E41L1LT, Percival Scientific, Inc., Perry, IA, USA) kept at 0 °C; other such three plug flats were transferred to another identical freezing chamber. The two freezing chambers, respectively, were used for freezing treatments of −5.5 or −6.5 °C, and subsequent thawing. The two test temperature treatments (−5.5 and −6.5 °C) used in the present study were selected based on our previous data of leaf-freezing response curve for ‘Reflect’ leaves, and LT_50_ (lethal temperature for 50 % injury) of ∼ −6.0 °C (Shin, Min, & Arora, 2018). These two test temperatures represent relatively moderate (−5.5 °C) and severe (−6.5 °C) stress bordering LT_50_, and are, therefore, physiologically relevant.

After 2-h at 0 °C, temperature in freezing chambers were lowed at 1 °C/h^-1^ up to -2 °C at which ice-nucleation was conducted by quickly misting pre-chilled (0°C) ddH_2_O onto leaves, and held at this temperature for 1-h. Plants were then frozen to −5.5 or −6.5 °C at -0.5 °C/30 min. Plants kept at each targeted temperature for 30 min were allowed to thaw at 0 °C overnight (∼13-h). Unfrozen control (UFC) seedlings of each treatment were kept at 0 °C in another identical chamber throughout the freeze-thaw cycle. Gradual thaw continued by subjecting plants, including UFC, to 5 °C for 2-h. Entire freezing and thawing was performed in dark. All the plants were transferred from chambers to the lab bench (∼20 °C) under dim light (∼15 μmol m^-2^s^-1^, cool white fluorescent) for ∼ 1-d. Freeze-thaw injury to seedlings was then evaluated visually and photographed. Additional assessment of freeze-injury / tolerance for F-control *versus* AsA-fed plants was made by measuring ion-leakage on leaves excised from plants that had been exposed to whole-plant freezing. Percent injury was calculated using percent ion-leakage data as described by Lim, Arora, & Townsend (1998).

Whole-plant freezing test along with visual estimation and ion-leakage measurement were repeated thrice, each with 14 to 16 plants per temperature per treatment (2 leaves per plant replicate). Injury percent data across three independent experiments were pooled to calculate the representative treatment means with standard errors. Mean differences were analyzed by LSD test.

#### 2.3.2 Excised-leaf freezing test (Bath freezing)

Excised-leaves from F-control and AsA-fed seedlings were subjected to a temperature-controlled freeze-thaw protocol as described by Chen & Arora (2014), using a glycol bath (Isotemp 3028; Fisher Scientific, Pittsburgh, PA, USA) (hereon referred as to ‘bath freezing’). Briefly, a pair of petiolate leaves (rinsed with ddH_2_O and blotted on paper towel) was placed in a 2.5 x 20 cm test tube containing 150 μl ddH_2_O and slowly cooled down at -0.5 °C/30min to four different test temperatures (i.e. -4.5, −5.5, −6.5, and −7.5 °C) following ice-nucleation at -1 °C. Samples were kept for 30 min at each selected temperature and thawed on ice overnight. UFC leaves of each treatment were maintained at 0 °C throughout the freeze-thaw cycle. The next morning, samples were kept at 4 °C for 1-h followed by 1-h at room temperature (∼20 °C) before measuring ion-leakage. Bath-freezing was independently repeated thrice, each with 5 technical replicates per temperature per treatment (2 leaves per technical replicate). Injury percent data (calculated from percent ion-leakage) from 3 biological replications were pooled to calculate the treatment means with standard errors. Mean differences were analyzed by LSD test.

### 2.4 ROS staining

Superoxide (O_2_ ^•−^) and hydrogen peroxide (H_2_O_2_) distribution was visualized by nitroblue tetrazolium (NBT) and 3,3’-diaminobenzidine (DAB) staining, respectively, using the protocol previously used in our laboratory for spinach (Chen & Arora, 2014; Min, Chen, & Arora, 2014). Staining intensities were visually evaluated between F-control and AsA-fed leaves that were subjected to bath-freezing at −5.5, −6.5, and −7.5 °C. This experiment was independently repeated twice, each with 2 to 3 replications (2 leaves / replicate) per temperature per treatment. A representative picture showing staining intensities is presented in this study.

### 2.5 Measurement of antioxidant enzyme activity

The activity of three antioxidant enzymes, i.e., SOD, CAT, and APX, was measured using a protocol as described by Chen & Arora (2014). Essentially, ground frozen leaf-tissue (150 mg) were homogenized with 1 ml of 100 mM potassium phosphate buffer (pH 7.0). The samples were then centrifuged at 10,000 *g* for 25 min at 4 °C and supernatants were used as the enzyme extract for SOD, CAT, and APX. Enzyme activity was calculated as described by Chen & Arora (2014) and Govinda, Sharma, Singh, & Jyoti (2017). This experiment was independently repeated four times, each with 3 to 4 technical replicates per treatment. Mean difference was analyzed as per Student’s *t*-test.

### 2.6 Measurement of glutathione (GSH)

GSH level was determined using high-performance liquid chromatography as described by Zheng et al. (2018) with slight modifications. Ground frozen leaf-tissues (∼ 0.2 g) were mixed with extraction buffer containing 0.1 % trifluoroacetic acid and 200 mM dithiothreitol to extract GSH. The homogenate was centrifuged at 15,300 *g* for 10 min. The supernatant (0.5 mL) was transferred to a spin filter and centrifuged for 5 min. The filtrate was injected into Spherisorb 5μm ODS column (250 mm x 4.6 mm) for HPLC (model 1260) coupled to 1200 series evaporative light scattering detector (Agilent Technologies, Santa Clara, CA, USA). This analysis was conducted twice independently with 2 to 3 technical replications each. Mean difference was analyzed as per Student’s *t*-test.

### 2.7 Sample extraction

Frozen leaf tissues were ground and used for metabolite profiling. Sample extraction was conducted as detailed by Min, Showman, Perera, & Arora (2018); each treatment from F-control and AsA-fed leaves consisted of 4 biological replications, each with 3 technical replications.

#### 2.7.1 Metabolite identification and quantification

Metabolite identification was performed based on compounds’ chromatographic retention time indices following deconvolution of raw GC-MS chromatograms using AMDIS software, as described by Min, Showman, Perera, & Arora (2018). Each identified metabolite was quantified based on internal standards and dry weight; missing data were replaced by a number (i.e. the smallest peak area /2) for further statistical analysis as reported by Xia, Psychogios, Young, & Wishart (2009).

#### 2.7.2 Statistical analysis for metabolite profiling

Principal component analysis (PCA) was conducted with R (version 3.2.2, The R Foundation for Statistical Computing, ISBN 3-900051-07-0) on log_10_ transformed relative metabolite concentration between two treatments (F-control *vs*. 1.0 mM AsA-fed tissues). Mean difference in the abundance of each metabolite between treatments was determined via Student’s *t*-test (Supplemental Table 1). A volcano plot was generated using log_2_-scaled mean difference in each metabolite concentration and log_10_-transformed p-values between two treatments; only those metabolites were numbered on a volcano plot for which the abundance between the two treatments was significantly different (p < 0.05).

### Supplementary Data

**Table S1.** Quantification of 46 metabolites with *t*-test between 2 biological conditions (F-control and 1.0 mM AsA).

**Table S2.** Loading values for 46 metabolites separated by two PCA components.

## 3 RESULTS

### 3.1 Effect of exogenous AsA on growth

Water content was slightly higher in 0.5 and 1.0 mM AsA-fed leaves compared to F-control, (Table 1). Leaf area of seedlings treated with 0.5 or 1.0 mM AsA was larger than the F-control by 7.0 or 15.9 %, respectively. DW/leaf area of F-control and 0.5 mM AsA-fed leaves was similar but slightly smaller than 1.0 mM AsA-fed seedlings.

**Table 1.** Leaf growth parameters of spinach (*Spinacia oleracea* L. cv. Reflect) seedlings sub-fertigated with fertilizer alone (F-control), fertilizer + 0.5 mM ascorbic acid (0.5 mM AsA), or fertilizer + 1.0 mM ascorbic acid (1.0 mM AsA). DW, dry weight.

| Growth parameters | Treatment |  |  |
| --- | --- | --- | --- |
|  | F-control | 0.5 mM AsA | 1.0 mM AsA |
| Water content (%) <sup>a</sup> | 91.9 ± 0.14 b | 92.5 ± 0.08 a | 92.3 ± 0.05 a |
| Leaf area (cm<sup>2</sup>) <sup>a</sup> | 6.30 ± 0.28 b | 6.74 ± 0.19 ab | 7.30 ± 0.29 a |
| DW/Leaf area (mg/cm<sup>2</sup>) <sup>a</sup> | 1.89 ± 0.03 b | 1.86 ± 0.03 b | 2.06 ± 0.02 a |

<sup>a</sup> Pooled means ± SE from five biological replications, each including 7–12 plants. Two leaves / plant were used resulting in a total of 14–24 leaves/biological replication. Values with the same letter within the same row are not different (LSD test, $p < 0.05$ )

### 3.2 Freezing tolerance and leaf-AsA

A representative picture of seedlings exposed to freeze-thaw stress is shown in Figure 1a where either 0.5 mM- or 1.0 mM-AsA fed seedlings are visually more freeze-tolerant than F-control at both −5.5 and −6.5 °C stress. The beneficial effect of AsA on FT was especially more pronounced at the moderate stress level (−5.5 °C). AsA (1.0 mM)-fed leaves accumulated ∼2.4-fold AsA compared to F-control (Figure 1b); leaf AsA of 0.5 mM AsA-fed leaves was not determined.

**Figure. 1.**
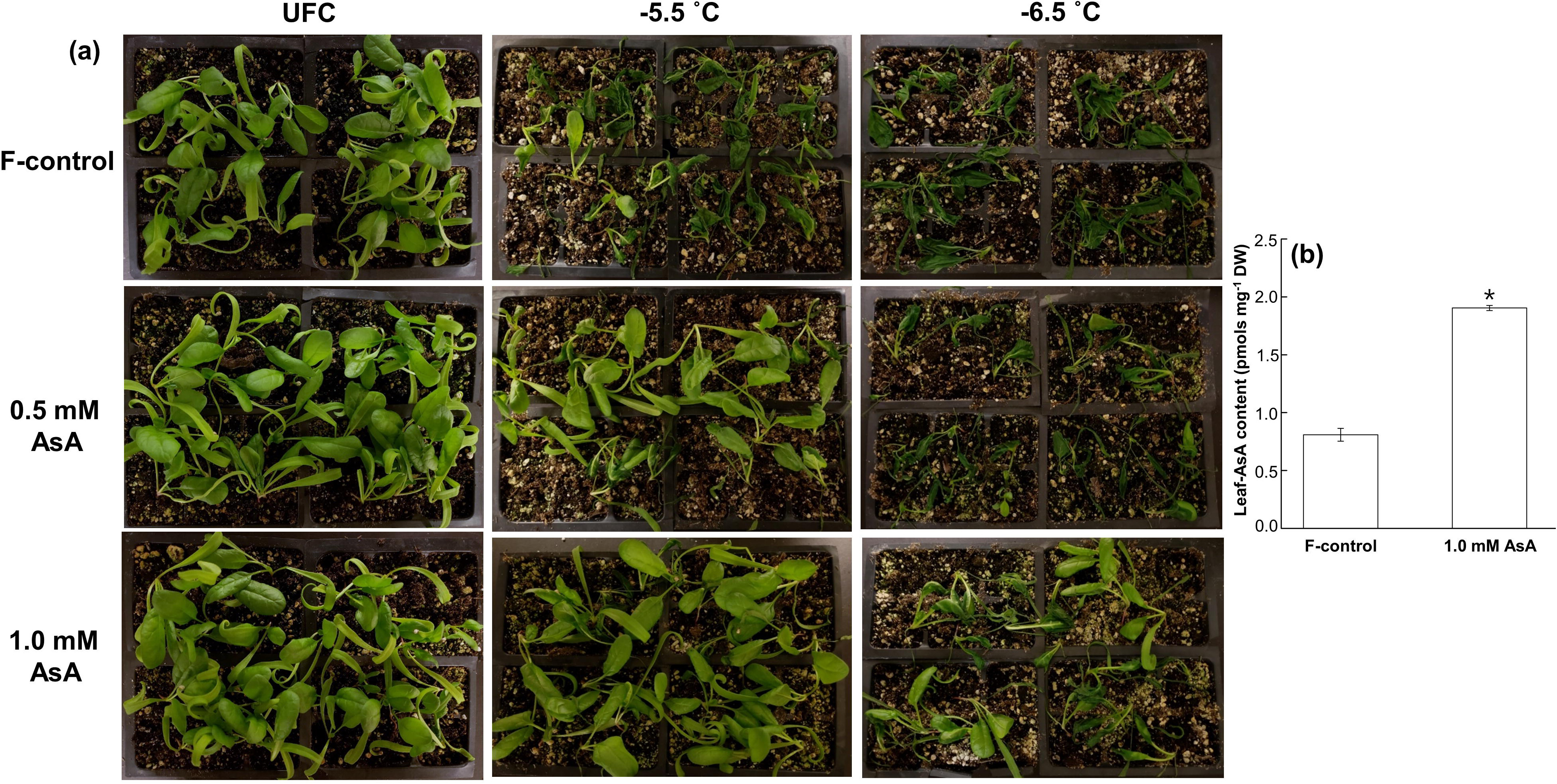
**(a)** Visual estimation of *in situ* whole-plant freezing (−5.5 and −6.5 °C) response of spinach (*Spinacia oleracea* L. cv. Reflect) seedlings sub-fertigated with fertilizer alone (F-control), fertilizer + 0.5 mM ascorbic acid (0.5 mM AsA), and fertilizer + 1.0 mM ascorbic acid (1.0 mM AsA); UFC, unfrozen control. **(b)** Ascorbic acid content (mean ± S.E.) in F-control and 1.0 mM AsA; different letters indicate significant differences between treatments at p < 0.05 as per LSD test.

Relatively less freeze-thaw injury in AsA-fed tissues was also evident by the ion-leakage from the leaves excised from seedlings that had been subjected to *in situ* freeze-thaw (Figure 2a). Seedlings fed with 0.5 mM AsA had ∼52 and ∼13% less injury at −5.5 and −6.5 °C, respectively, compared to F-control whereas those treated with 1.0 mM AsA had ∼ 69 and ∼ 41% less injury at both stress levels relative to F-control. Bath-freezing tests using excised leaves (not whole seedlings) from three different treatments also exhibited lower freezing injury in AsA-fed tissues compared to F-control (Figure 2b).

**Figure. 2.**
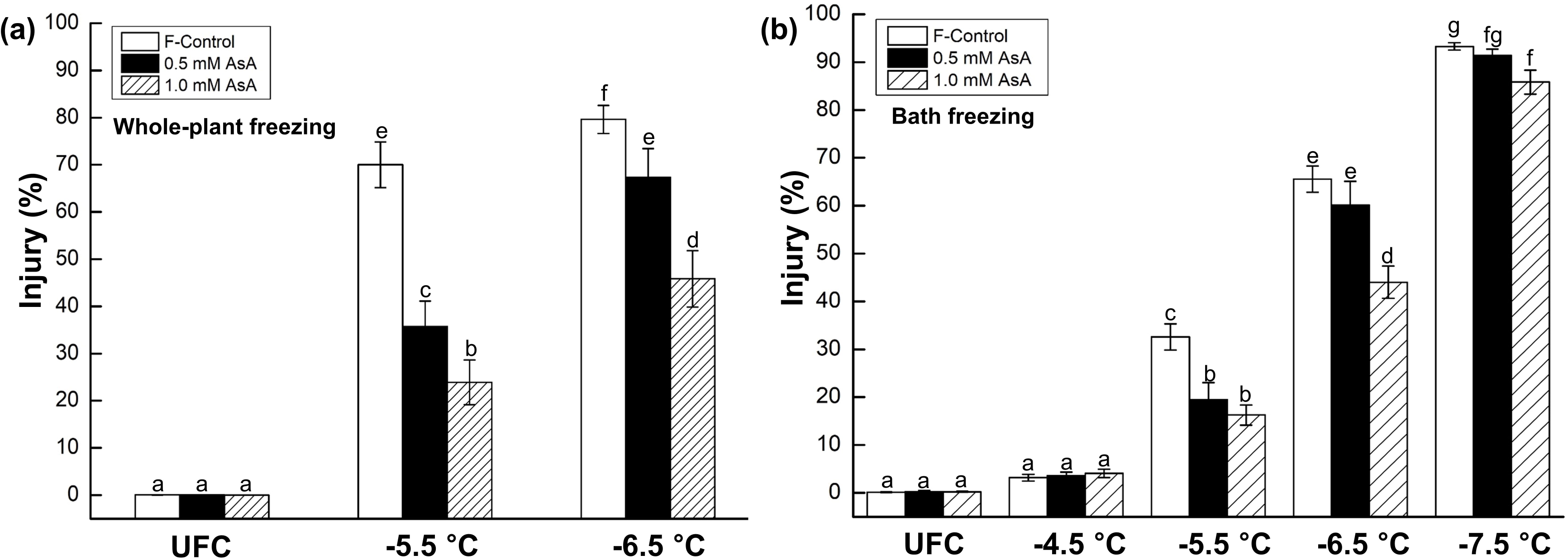
Freeze-thaw injury assessed by ion-leakage of spinach (*Spinacia oleracea* L. cv. Reflect) leaves: **(a)** excised from the seedlings subjected to *in situ* whole-plant freezing at −5.5 and −6.5 °C; values represent the average ± S.E. from three independent experiments, each with 14-16 plants per temperature per treatment and **(b)** exposed to bath-freezing at -4.5, −5.5, −6.5, and −7.5 °C; values represent the average ± S.E. from three independent experiments, each with 5 plants per temperature per treatment. Different letters indicate significant differences between treatments at p < 0.05 as per LSD test. UFC, unfrozen control; F-control, seedlings sub-fertigated with fertilizer alone; 0.5 mM AsA, seedlings sub-fertigated with fertilizer + 0.5 mM ascorbic acid; 1.0 mM AsA, seedlings sub-fertigated with fertilizer + 1.0 mM ascorbic acid.

### 3.3 Histochemical detection of ROS (O_2_ ^•−^ and H_2_O_2_)

A representative image of the quantitative estimate of O_2_ ^•−^ and H_2_O_2_ (as indicated by the color intensity) in the leaves from three treatments (F-control, 0.5 mM and 1.0 mM AsA) after having been exposed to bath-freezing at −5.5, −6.5, and −7.5 °C, and that of unfrozen control (UFC) is shown in Figure 3. The two ROS accumulated at higher abundance in F-control than 0.5 and 1.0 mM AsA-fed leaves after freezing at −5.5 and −6.5 °C, with 1.0 mM AsA treatment showing the lowest accumulation. Little to no protection was apparent by AsA application at −7.5 °C stress level.

**Figure. 3.**
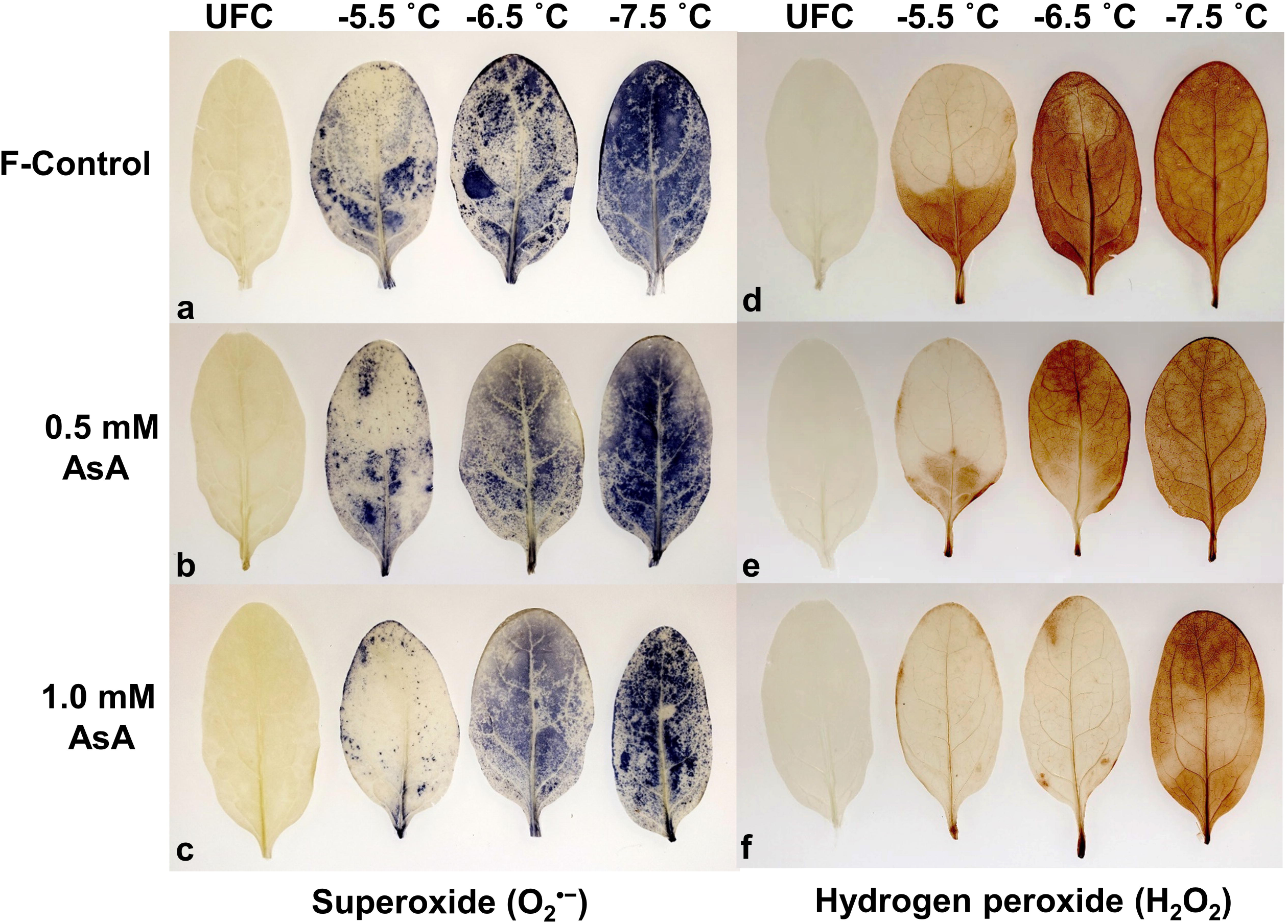
Distribution of superoxide (O_2_ ^•−^) **(a-c)** and hydrogen peroxide (H_2_O_2_) **(d-f)** in unfrozen controls (UFC) and freeze-thaw injured spinach (*Spinacia oleracea* L. cv. Reflect) leaves that were sub-fertigated with fertilizer alone (F-control), fertilizer + 0.5 mM ascorbic acid (0.5 mM AsA), and fertilizer + 1.0 mM ascorbic acid (1.0 mM AsA) before exposure to bath-freezing at −5.5, −6.5, and −7.5 °C.

### 3.4 Biochemical analysis

FT data indicated 1.0 mM AsA treatment to be more protective than 0.5 mM. Therefore, F-control was hereon compared only with 1.0 mM AsA treatment for all biochemical analyses (below).

### 3.5 Antioxidant enzyme activities and leaf-glutathione (GSH)

Quantification of antioxidant enzyme activities and GSH was expressed on DW basis, since water content of F-control *versus* AsA-fed leaves was different (Table 1).

SOD (Figure 4a), CAT (Figure 4b), and APX (Figure 4c) activities in 1.0 mM AsA-fed leaves, respectively, were 1.1-, 2.4- and 2.7-fold of F-control. GSH in 1.0 mM AsA-fed leaves was ∼1.3-fold of F-control (Figure 4d).

**Figure. 4.**
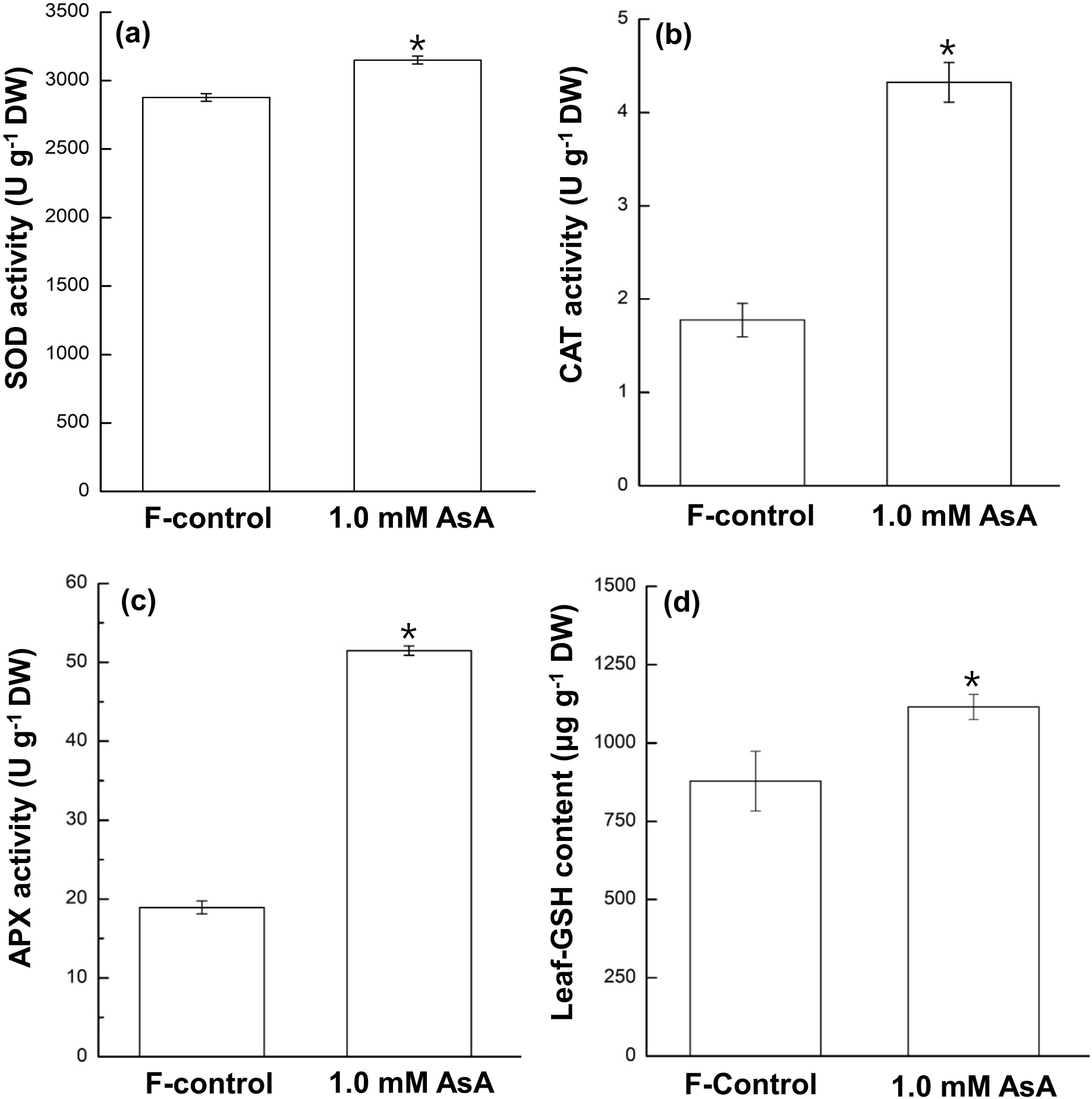
The activity of SOD, CAT and APX **(a-c)** in spinach (*Spinacia oleracea* L. cv. Reflect) leaves sub-fertigated with fertilizer alone (F-control), and fertilizer + 1.0 mM ascorbic acid (1.0 mM AsA). One unit of SOD activity is defined as the amount of enzyme required for 50 % inhibition of formazan formation at 560 nm; one unit of CAT activity is defined as the degradation of 1 μM H_2_O_2_ in 1 min at 240 nm; one unit of APX activity is defined as the degradation of 1 μM AsA into monodehydroascorbate in 1 min at 290 nm; values represent the average ± S.E from four independent experiments, each with 3 - 4 replications per treatment. **(d)** Glutathione concentration of spinach leaves in F-control and 1.0 mM AsA; values represent the average ± S.E from two independent experiments, each with 2 – 3 replications per treatment. * indicates significant difference at p < 0.05 (*t*-test) for all four panels.

### 3.6 Principal Component analysis (PCA)

In total, 46 metabolites were identified following GC-MS analysis and clustered into 6 groups— 17 amino acids, 8 carbohydrates, 2 fatty acids, 4 TCA intermediates, 3 antioxidants and 12 others (Supplemental Table 1).

PCA was performed to explore whether metabolite phenotype between F-control and 1.0 mM AsA were different, and to determine which metabolites affected such differences the most. Data indicated clear separation between the two treatments wherein two components (PC1 and PC2) explained 55.6 % of the total variance (Figure 5a). The first component (PC1) accounting for 37.4 % of the variance separating AsA-feeding *versus* F-control. The second component (PC2) accounting for 18.2 % of the variance primarily indicate different abundance of metabolites across technical replications within each treatment (Figure 5a). Loading values for metabolites separated by the two PCs are shown in Supplemental Table 2. For example, urea and GABA with most positive or negative loading values, respectively, contribute most for the separation of F-control against AsA-fed treatment on PC1.

**Figure. 5.**
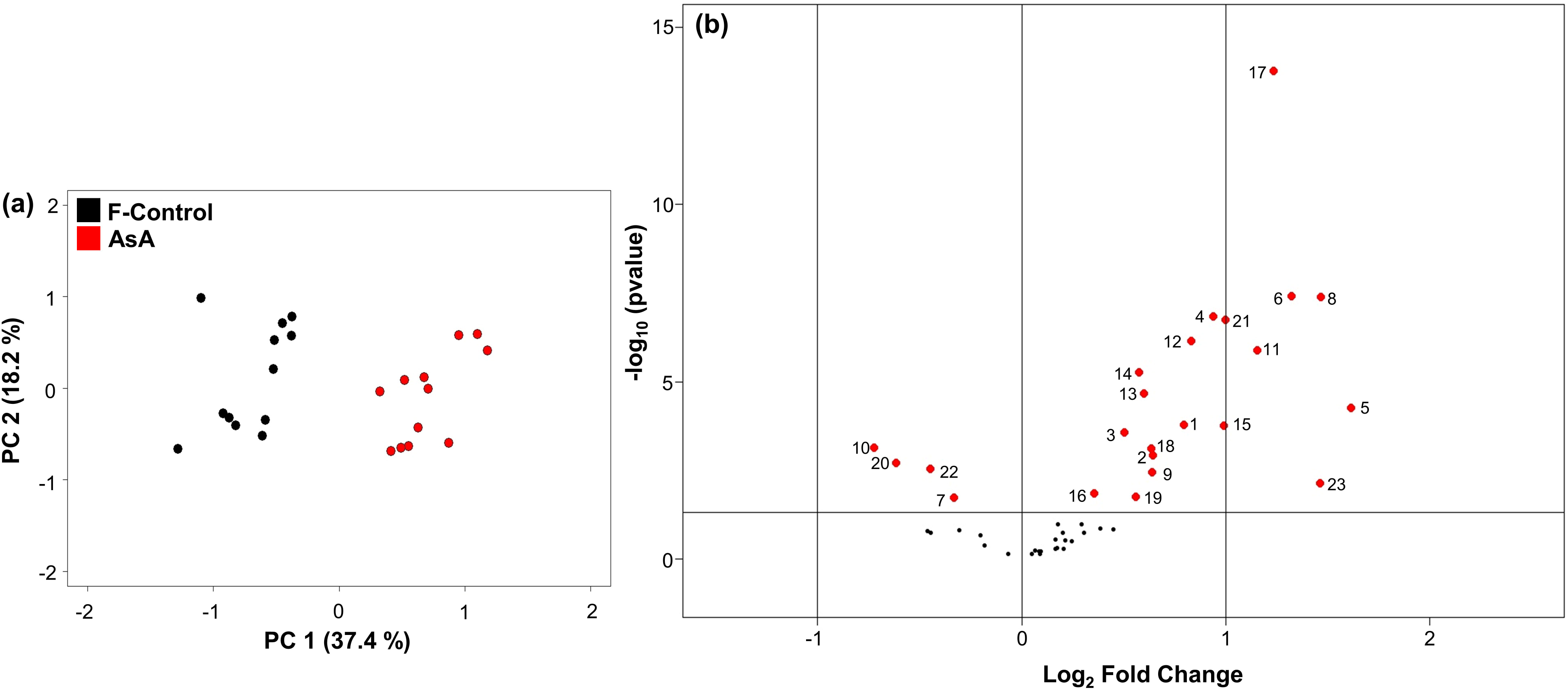
**(a)** Metabolic phenotype clustering for spinach (*Spinacia oleracea* L. cv. Reflect) leaves through principal component analysis (PCA) of log_10_-scaled 46 metabolite data from a total of 12 replications (triplicates from 4 biological replications) each originating from two treatments (i.e. F-control and 1.0 mM AsA); 1.0 mM AsA: treated with fertilizer + 1.0 mM AsA; F-control: treated with fertilizer. Principal component 1 (PC1) indicates differential response to AsA application. Principal component 2 (PC2) indicates variation of metabolite concentration between replications. F-control (black) and 1.0 mM AsA (red) are shown in 2D plot. **(b)** Volcano plot of comparative abundance of metabolites in 1.0 mM AsA *vs.* F-control. Each dot represents a metabolite with the -log_10_ (p value) as a function of abundance difference between two biological conditions (log_2_ fold change on the abscissa). Metabolites are numbered and colored (red) if significantly different at p < 0.05. The two vertical lines on either side of the central vertical line indicate range of two-fold cut off in abundance whereas the horizontal line represents a threshold of -log_10_ = 0.05.

### 3.7 Comparative metabolite profiles of 1.0 mM AsA vs. F-control

Mean abundance of total 46 identified metabolites was pair-wise compared (1.0 mM AsA *vs*. F-control) using log_2_-scaled fold change and -log_10_ scaled p-values. Data indicate that 23 out of 46 metabolites marked as red in Figure 5b exhibited major changes in abundance (significantly different at p < 0.05) between F-control and AsA-fed treatment. Numerical fold-change (log_2_ scaled) for these metabolites in AsA-fed *vs.* F-control are shown in Table 2, where the spot number for each metabolite corresponds to the number assigned in Figure 5b. These 23 metabolites are also identified for significance level (*t*-test) with asterisk notations in Supplemental Table 1. Nineteen of these metabolites, i.e. cysteine, glycine, glutamine, glutamic acid, leucine, methionine, proline, threonine, galactinol, myo-inositol, citric acid, malic acid, α-tocopherol, γ-tocopherol, AsA, ferulic acid, glyceric acid, phytol, and urea, were more abundant in 1.0 mM AsA-fed leaves as indicated by a positive value (1.0 mM AsA/ F-control ratio) (Table 2). Four metabolites, phenylalanine, fructose, GABA, and phosphoric acid were less abundant (minus sign) relative to F-control (Table 2). These 23 metabolites were placed in five categories (not 6, as in Supplemental Table 1) because abundance in ‘fatty acids’ was not found to be significantly different.

**Table 2.**
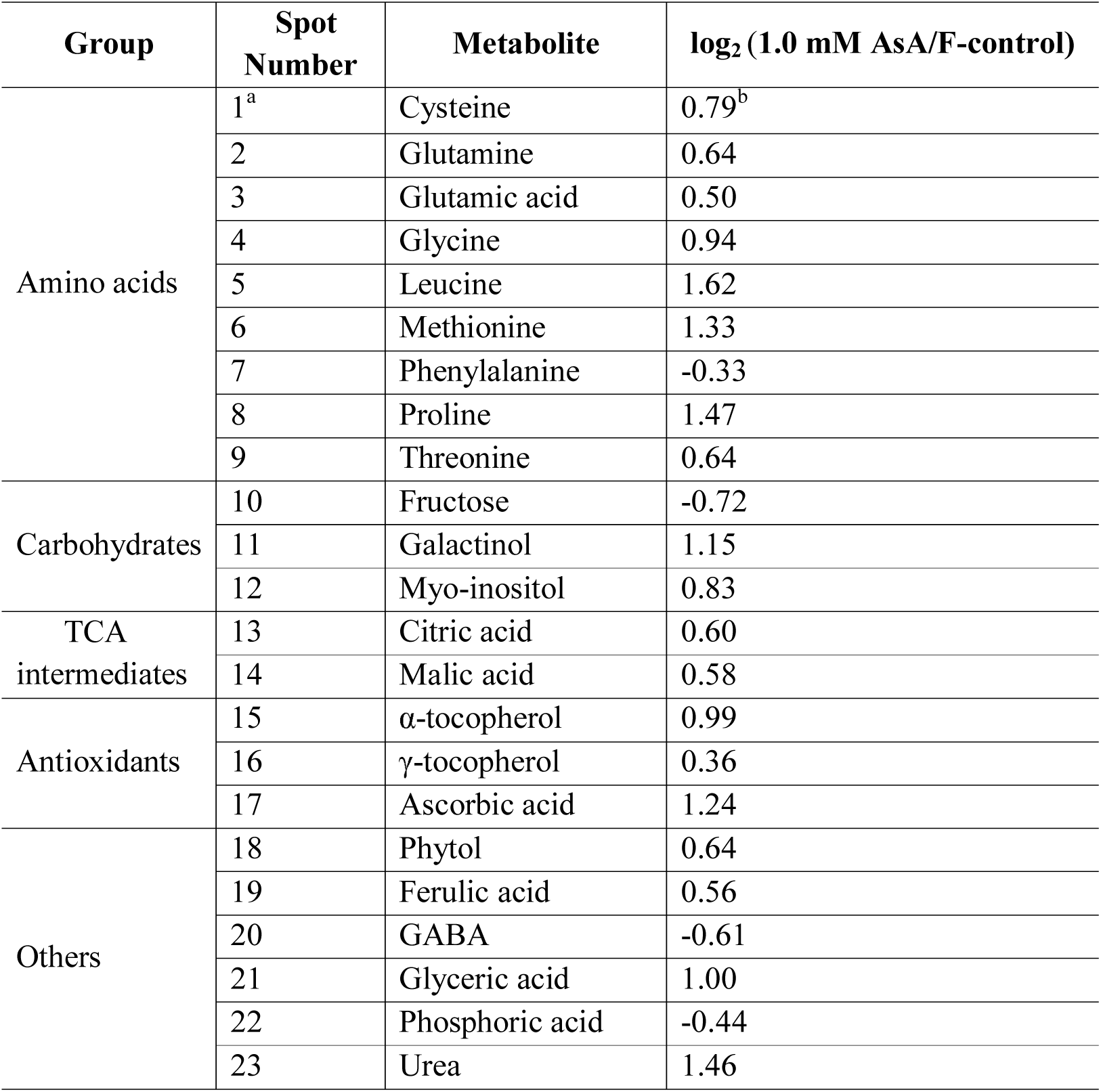
Significant changes in the concentrations of leaf-metabolites between 1.0 mM AsA *vs.* F-control. ^a^ spot number indicates a metabolite that is significantly different (p < 0.05) and ^b^ fold changes in the concentrations of each metabolites between two groups (12 replications per treatment) were calculated using the formula log_2_ (1.0 mM AsA / F-control); spot numbers and the numerical value of metabolites in this table are illustrated in Fig. 5b. Metabolites are classified into 5 groups, i.e. amino acids, carbohydrates, TCA intermediates, antioxidants, and others. 1.0 mM AsA: seedlings sub-fertigated with fertilizer + 1.0 mM ascorbic acid; F-control: seedlings sub-fertigated with fertilizer alone.

| Group | Spot Number | Metabolite | $\log_2 (1.0 \text{ mM AsA/F-control})$ |
| --- | --- | --- | --- |
| Amino acids | 1<sup>a</sup> | Cysteine | 0.79<sup>b</sup> |
|  | 2 | Glutamine | 0.64 |
|  | 3 | Glutamic acid | 0.50 |
|  | 4 | Glycine | 0.94 |
|  | 5 | Leucine | 1.62 |
|  | 6 | Methionine | 1.33 |
|  | 7 | Phenylalanine | -0.33 |
|  | 8 | Proline | 1.47 |
|  | 9 | Threonine | 0.64 |
| Carbohydrates | 10 | Fructose | -0.72 |
|  | 11 | Galactinol | 1.15 |
|  | 12 | Myo-inositol | 0.83 |
| TCA intermediates | 13 | Citric acid | 0.60 |
|  | 14 | Malic acid | 0.58 |
| Antioxidants | 15 | α-tocopherol | 0.99 |
| | 16 | γ-tocopherol | 0.36 |
|  | 17 | Ascorbic acid | 1.24 |
| Others | 18 | Phytol | 0.64 |
|  | 19 | Ferulic acid | 0.56 |
|  | 20 | GABA | -0.61 |
|  | 21 | Glyceric acid | 1.00 |
|  | 22 | Phosphoric acid | -0.44 |
|  | 23 | Urea | 1.46 |

## 4 DISCUSSION

In recent years, exogenous application of beneficial chemicals has received some attention as potential means for improving plant tolerance against various abiotic stresses (Savvides, Ali, Tester, & Fotopoulos, 2016). While AsA application has been a subject of such efforts in the context of salt, drought and chilling stresses, its effect on FT remains unknown. In the present study, we have evaluated the effect of AsA-fertigation on FT at whole-plant as well as excised tissue level determined through various parameters of freeze-thaw injury, and conducted metabolite profiling of leaves before and after AsA-treatment to explore metabolic explanation for AsA-mediated change in FT.

### 4.1 AsA-fertigation and leaf-growth

The effect of AsA on plant growth as well as stress tolerance is dependent upon the mode of application and concentration (Akram, Shafiq, & Ashraf, 2017). Hence, we first tested four AsA concentrations (i.e. 0.5, 1.0, 2.0, and 4.0 mM) as sub-fertigation treatments. Seedlings fed with 2.0 and 4.0 mM AsA showed somewhat stunted growth relative to F-control, whereas 0.5 and 1.0 mM AsA feeding did not show any detrimental effect. Hence, 2.0 and 4.0 mM AsA were not used for subsequent experiments (data not shown for these comparisons). Higher leaf AsA concentration in AsA-fed leaves than F-control (Figure 1b) indicates seedlings effectively absorbed and assimilated AsA.

Research shows that exogenous application of AsA improved plant growth in wheat (Athar, Khan, & Ashraf, 2008) and millet (Hussein & Alva, 2014). This supports our observation of 7-16% higher leaf-area in AsA-fed leaves (Table 1). Specific mechanism of AsA-induced growth is beyond the scope of this study but increase in AsA level has been associated with enhanced cell division (Smirnoff, 1996) and expansion (De Cabo, González-Reyes, Córdoba, & Navas, 1996). Moreover, repression of _L_-galactono-1,4-lactone dehydrogenase, an enzyme involved in the biosynthesis of AsA, in tobacco BY-2 cell lines caused a decline in cellular AsA content as well as in cell division and growth (Horemans, Potters, Wilde, & Caubergs, 2003). In contrast, cell division in maize root treated with AsA was higher than control (Kerk & Feldman, 1995). AsA may promote cell division by inducing G_1_ to S progression (Galie, 2013) and AsA-induced plant growth could also involve auxin regulation via interaction between ascorbate oxidase and auxin (Key, 1962; Esaka, Fujisawa, Goto, & Kisu, 1992; Smirnoff, 2000; Pignocchi, Fletcher, Wilkinson, Barnes, & Foyer, 2003).

### 4.2 AsA-feeding improves freezing tolerance

Visual evaluation of injured seedlings (whole plant freezing) and corresponding percent injury based on ion-leakage from leaves excised from these seedlings indicate AsA-fed plants to be more freeze-tolerant than F-control (Figure 1a and 2a), and that 1.0 mM AsA was more effective than 0.5 mM AsA. ‘Bath-freezing’ tests with excised spinach leaves further supported this observation (Figure 2b). Induction of freezing tolerance (as in cold acclimation) typically involves decrease in cellular hydration status (Xin & Browse, 2000). Therefore, it is somewhat intriguing that AsA-fed leaves, which are more hydrated, though marginally, than F-control (Table 1), had greater FT. Conceivably, other physiological and biochemical changes induced by AsA-feeding (as discussed below) override this apparent contradiction.

### 4.3 Higher antioxidant enzyme activity in AsA-fed leaves

Evidence abounds that plant tissues subjected to freeze-thaw accumulate excess O_2_ ^•−^ and H_2_O_2_ (Kendall and McKersie, 1989; Min, Chen, & Arora, 2014; Shin, Min, & Arora, 2018). Our data of visual detection of ROS (Figure 3) is consistent with these findings and shows that more severe freezing stress (−6.5 °C) resulted in higher ROS accumulation. More importantly, less accumulation of O_2_ ^•−^ and H_2_O_2_ in AsA-fed leaves compared to F-control indicates alleviation of oxidative stress by AsA application. Relatively higher scavenging of H_2_O_2_ in 1.0 mM AsA-feeding compared to 0.5 mM (compare Figure 3e and f) indicates the former treatment was more effective scavenger of free radicals. Our results also indicate a close correspondence between antioxidant enzyme activities and ROS abundance. For instance, SOD activity in AsA-fed leaves was higher than F-control (Figure 4a), which might be responsible for less accumulation of O_2_ ^•−^. Likewise, CAT and APX activities in AsA-fed leaves were higher than in F-control (Figure 4b, c), suggesting more efficient scavenging of H_2_O_2_. A higher GSH content in 1.0 mM AsA-fed leaves than F-control (Figure 4d) further supports higher APX activity in these tissues since GSH works together with APX in ascorbate-glutathione cycle (Foyer & Noctor, 2011). Several studies have also noted enhanced activity of antioxidant enzymes by exogenous application of AsA, especially when tissues are exposed to abiotic stresses (Athar, Khan, & Ashraf, 2008; Kumar et al., 2011; Alam, Nahar, Hasanuzzaman, & Fujita, 2014).

### 4.4 AsA-feeding alters leaf metabolome

PC1, accounting for 37.4% of total variance, clearly separated F-control from AsA-fed treatment (Figure 5a). In contrast, PC2, explaining 18.2% of total variance, may reflect differences of metabolite concentration across replications. Discussion under several sections (below) further highlights specific differences in metabolism between two treatments.

Figure 5b illustrates comparative metabolite abundance for AsA-fed *vs.* F-control tissues. Ensuing metabolite-specific sections discuss their putative roles in relation to FT.

#### 4.4.1 Amino acids

AsA-fed leaves had significantly higher levels of cysteine, methionine, proline, glutamine, glutamic acid, glycine, threonine and leucine but lower level of phenylalanine relative to F-control (spots 1-9; Figure 5b; Table 2). Higher cysteine, glutamic acid, and glycine in AsA-fed tissues supports our results of a higher activity of APX as well as higher GSH in these tissues (Figure 4c, d). Cysteine, a sulfur-containing amino acid, is known as a key component for GSH biosynthesis (Noctor et al., 2012), which consists of two steps: 1) formation of γ-glutamyl-cysteine, catalyzed by glutamate-cysteine ligase, and 2) addition of glycine (or, β-alanine, serine and glutamic acid) to γ-glutamyl-cysteine, catalyzed by glutathione synthase. GSH is involved in AsA-GSH cycle as a substrate for dehydroascorbate reductase which reduces dehydroascorbate to ascorbate (Smirnoff, 2000; Foyer & Noctor, 2011). Others have also noted a higher accumulation of AsA and GSH induced by exogenous AsA under heat (Kumar et al., 2011) and salt stress (Billah, Rohman, Hossain, & Uddin, 2017).

Methionine is an indispensable building block for protein synthesis. Higher methionine in AsA-fed leaves may be useful for the synthesis of various stress-proteins associated with FT induction (Espevig, Xu, Aamlid, DaCosta, & Huang, 2012; Chen et al., 2015). Arrigoni, Arrigoni-Liso, & Calabrese (1977) reported that AsA was necessary to synthesize hydroxyproline-containing proteins, a cell wall structural entity important for cell expansion / growth (Cleland, 1968; Ridge & Osborne, 1971; Kavi Kishor, Hima Kumari, Sunlta, & Sreenivasulu, 2015). Conceivably, higher methionine in AsA-fed leaves may also be associated with small but significantly better leaf-growth of AsA-fed seedlings (Table 1). Methionine also serves as a substrate for the synthesis of polyamines (Alcázar et al., 2011). Accumulation of polyamines has been implicated in stress tolerance, including freezing (Alcázar et al., 2011).

Phenylalanine, an aromatic amino acid, serves as a precursor for a wide range of important secondary metabolites (Tzin & Galili, 2010). One such metabolite, lignin, a strengthening component of cell wall, is synthesized via phenylpropanoid/lignin biosynthetic pathway (Vanholme, Demedts, Morreel, Ralph, & Boerjan, 2010). In the present study, AsA-fed leaves had lower levels of phenylalanine. This may be due to either decreased synthesis or increased consumption of phenylalanine, the latter presumably for lignin biosynthesis.

Lignin content was not measured in this study. However, higher lignin content has been widely linked with increased FT (Huner, Palta, Li, & Carter, 1981; Stefanowska, Kuras, Kubacka-zebalska, & Kacperska, 1999). Cold acclimation-induced upregulation of C3H gene (a key enzyme for lignin biosynthesis) has also been reported for *Rhododendron* leaves (Wei et al., 2006). Higher lignin content is also expected with greater tissue growth as well as higher leaf DW; higher leaf area and DW/leaf area for AsA-fed tissues in this study is in line with this notion. Future study of lignin biosynthesis and content in AsA-fed tissues is warranted to test above stated notion.

AsA-fed leaves had ∼2.8-fold proline relative to F-control (spot 8; Figure 5b; Supplemental Table 1). Proline, a compatible solute, has been widely known to accumulate under stress conditions with roles in cellular osmotic adjustment, and membrane and protein stabilization (Hayat et al., 2012). Its accumulation has also been widely reported in cold-acclimated plants including spinach (Kaplan et al., 2004; Shin, Min, & Arora, 2018; Min, Showman, Perera, & Arora, 2018). Concordantly, AsA-fed plants were also more freeze-tolerant in the present study. How AsA-feeding causes proline accumulation is not known. However, a relatively higher amount of glutamine and glutamic acid in AsA-fed leaves compared to F-control (spot 2, 3; Figure 5b; Table 2) suggests stimulation of proline biosynthesis since glutamine is converted into glutamic acid, a primary precursor of proline biosynthesis (Forde and Lea, 2007; Hayat et al., 2012). Indeed, AsA-induced proline accumulation has been reported in okra under drought (Amin, Mahleghah, Mahmood, & Hossein, 2009) and wheat under salt stress (Azzedine, Gherroucha, & Baka, 2011). On the other hand, Hoque et al. (2007) reported that activity of enzymes involved in AsA-GSH cycle, including APX, were stimulated by exogenous proline in tobacco cultures under salt stress.

Leucine, a branched amino acid, was significantly higher in AsA-fed leaves (spot 5; Figure 5b; Table 2). Although not significantly, other branched amino acids, isoleucine and valine, also were higher in these tissues compared to F-control (Supplemental Table 1). Branched amino acids serve as precursors for the biosynthesis of secondary metabolites involved in various plant defenses (Bennett & Wallsgrove, 1994; Dixon, 2001). Also, upregulation of genes involved in secondary metabolism have been well correlated with improved FT (Hannah et al., 2006). Hence, higher abundance of branched amino acids in AsA-fed leaves may indicate higher level of secondary metabolites specifically contributing to higher FT. However, this notion deserves further confirmation.

Threonine (spot 9; Figure 5b; Table 2) was higher in AsA-fed leaves relative to F-control, but no explanation is available at this time for their role in FT induction.

#### 4.4.2 Carbohydrates

Fructose (as well as glucose) was less abundant in AsA-fed leaves compared to F-control (spot 10; Figure 5b; Table 2). Reason for this is unclear but may have resulted from decreased breakdown of sucrose which accumulated at higher levels in AsA-fed (Supplementary Table 1). Higher sucrose in AsA-fed leaves may be associated with increased FT due to its well established role as a compatible solute under desiccation stress (Kaplan et al., 2004; Bocian et al., 2015).

AsA-fed leaves had higher abundance of galactinol and myo-inositol (spot 11, 12; Figure 5b; Table 2). Galactinol and myo-inositol are involved in the biosynthesis of raffinose family oligosaccharides (RFOs) (Sengupta, Mukherjee, Basak, & Majumder, 2015; Kannan et al., 2016). RFOs also serve as compatible solutes under stress conditions (Kaplan et al., 2004; Bocian et al., 2015). Hincha, Zuther, & Heyer (2003) noted that RFOs stabilized cellular membrane under desiccation stress via sugar-membrane interaction. Although RFOs were not detected in the present study due to technical limitations (oligosaccharides were undetectable by GC-MS used here), it may be reasonable that AsA-fed leaves have higher abundance of RFOs which may contribute, in part, to enhanced FT.

#### 4.4.3 Antioxidants

Alpha-tocopherol and gamma-tocopherol were ∼2.0- and ∼1.3-fold of F-control, respectively (spot 15, 16; Figure 5b; Supplemental Table 1), indeed a substantially high accumulation. Alpha-tocopherol is a potent antioxidant protecting membranes by scavenging singlet oxygen and reacting with lipid peroxyl radicals, i.e. reduced lipid peroxidation (Sattler, Gilliland, Magallanes-Lundback, Pollard, & DellaPenna, 2004; Munné-Bosch, 2005), which may contribute to amelioration of freeze-injury. The exact mechanism of how AsA-feeding induces accumulation of alpha-tocopherol is unclear. However, data from the present study could provide tentative explanation as follows: tocopherol biosynthesis requires phytyl-diphosphate which is derived by phosphorylation of free phytol (Soll & Schultz, 1981; Vom Dorp et al., 2015). AsA-fed leaves, in the present study, had higher free phytol levels than F-control (spot 18; Figure 5b; Table 2). Moreover, it has been noted that tocopheroxyl radical, i.e. oxidized form of tocopherol, is reduced by ascorbic acid and therefore, tocopherols and AsA work collaboratively in controlling ROS levels (Munné-Bosch, 2005; Szarka, Tomasskovics & Bánhegyi, 2012). AsA and GSH (discussed earlier) in conjunction with alpha-tocopherol constitute a robust antioxidant machinery in AsA-fed leaves enabling greater resistance to freeze-induced oxidative stress.

#### 4.4.4 TCA intermediates and other metabolites

Citric acid and malic acid were more abundant in AsA-fed leaves compared to F-control (spots 13, 14; Figure 5b; Table 2); although not significantly different, two other TCA intermediates, fumaric acid and succinic acid, were also more abundant in AsA-fed leaves (Supplemental Table 1). TCA cycle is pivotal in producing energy for various biochemical processes and delivering carbon skeleton and reducing equivalents (Meyer et al., 2007). Hence, this bigger pool size of TCA metabolites in AsA-fed leaves may be associated with accumulation of many useful metabolites which contribute to improved FT.

GABA was higher in F-control compared to AsA-fed leaves (spot 20; Figure 5b; Table 2). GABA is a four-carbon non-proteinogenic amino acid requiring glutamic acid as a precursor for its synthesis (Shelp, Bown, & McLean, 1999). In the present study, AsA-fed leaves had substantially higher proline, which too requires glutamic acid for its biosynthesis. Could it be that lower level of GABA in AsA-fed tissues resulted from the lack of sufficient precursor? This hypothesis warrants further confirmation. Phosphoric acid was also higher in F-control than AsA-fed leaves (spot 22; Figure 5b; Table 2) whereas ferulic acid, glyceric acid, and urea were more abundant in AsA-fed leaves than F-control (spots 19, 21, 23; Figure 5b; Table 2). No explanation is available at this time for their specific role, if any, in FT.

## 5. CONCLUSION

Summarized conclusions are illustrated in **Figure 6**. AsA-feeding of spinach seedlings enhanced activity of SOD, CAT, and APX, and bolstered the accumulation of antioxidants (alpha- and gamma-tocopherol, AsA, glutathione) and compatible solutes/osmolytes (proline, galactinol and myo-inositol). These changes may synergistically enhance FT via alleviating freezing-induced oxidative stress as well as protect membranes from freeze-desiccation. Additional components of improved FT of AsA-fed leaves can be enhanced secondary metabolite system and lignin / cell wall augmentation; these two presumed changes are supported by increase in branched amino acids (leucine, isoleucine, valine) and possibly higher consumption of phenylalanine, respectively. Lastly, AsA-feeding induced small but significant increase in leaf-growth is possibly a result of enhanced expansion and/or division.

**Figure. 6.**
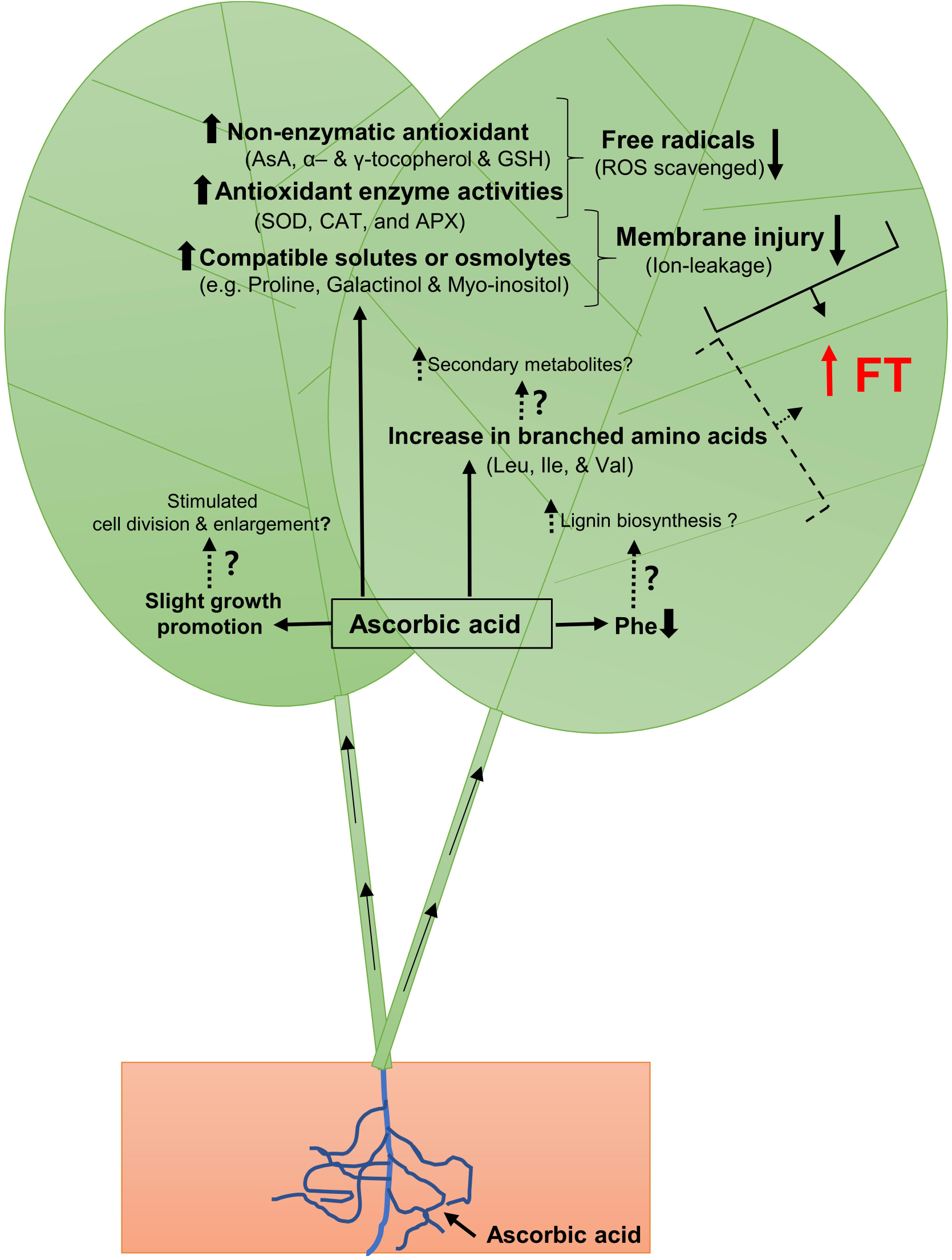
Illustrative summary of the effect of exogenous application of AsA on leaf metabolome *vis-à-vis* improved freezing tolerance (FT) of spinach (*Spinacia oleracea* L. cv. Reflect); for explanation, refer to ‘Conclusions’. AsA, ascorbic acid; GSH, glutathione; SOD, superoxide dismutase; CAT, catalase; APX, ascorbate peroxidase; ROS, reactive oxygen species; Leu, leucine; Ile, isoleucine; Val, valine; Phe, phenylalanine.

## ACKNOWLEDGEMENTS

This journal paper of the Iowa Agriculture and Home Economics Experiment Station, Ames, Iowa, Project no. 3601 was supported by Hatch Act and State of Iowa Funds. Technical assistance by Mr. Peter Lawlor (Manager, Horticulture Greenhouses) and Drs. Ann Perera and Lucas Showman (W.M. Keck Metabolomics Laboratory), Iowa State University is gratefully acknowledged.

## AUTHOR CONTRIBUTIONS

R.A. and K.M. jointly conceived the idea and designed experiments. K.M. performed the experiments and analyzed the data with help from K.C. K.M. and R.A. jointly wrote the paper. R.A. provided all financial support for this research.

