## Supplementary Table 1, 2 for "A metabolomics study of ascorbic acid-induced *in situ* freezing tolerance in spinach (*Spinacia oleracea* L.)"

**Supplementary Table 1.** Concentrations (p.mols mg^-1^ DW) of 46 metabolites in spinach leaves across 2 biological conditions (F-control and 1.0 mM AsA) as detected by GC-MS. Fresh weight (average of 12 replicates; triplicates from four biological replications) used for metabolite profiling experiments was 30.5 and 31.0 mg for F-control and 1.0 mM AsA, respectively. Dry weight in a given amount of fresh weight for F-control and 1.0 mM AsA, respectively, was 2.45 and 2.37 mg, as extrapolated using the percent water content from Table 1. Values are averages of 12 replications (n). Metabolites are grouped under 6 categories.

| **Categories** | **Metabolite** | **F-control**  **(n=12)** | **1,0 mM**  **AsA (n=12)** | **1.0mM/**  **F-control** | ***t*-test** |
| --- | --- | --- | --- | --- | --- |
| **Amino acids**  **(17)** | Beta-alanine | 0.04171 | 0.047055 | 1.1281467 | ns |
|  | 5-oxoproline | 2.049614 | 2.363335 | 1.1530635 | ns |
|  | Asparagine | 0.67263 | 0.48839 | 0.7260901 | ns |
|  | Glycine | 0.301301 | 0.578739 | 1.9208001 | **** |
|  | Cysteine | 0.113125 | 0.196312 | 1.7353547 | *** |
|  | Glutamic acid | 44.339 | 62.88653 | 1.4183119 | *** |
|  | Glutamine | 2.447678 | 3.823117 | 1.5619363 | ** |
|  | Isoleucine | 0.394166 | 0.538268 | 1.3655871 | ns |
|  | Leucine | 1.457205 | 4.469551 | 3.0672081 | **** |
|  | Methionine | 0.192582 | 0.482649 | 2.5062 | **** |
|  | Proline | 1.003127 | 2.777227 | 2.7685697 | **** |
|  | Serine | 18.66626 | 20.95382 | 1.1225505 | ns |
|  | Threonine | 1.982117 | 3.091374 | 1.5596325 | ** |
|  | Tyrosine | 1.178144 | 1.234539 | 1.0478677 | ns |
|  | Valine | 6.675684 | 7.668362 | 1.1487006 | ns |
|  | Aspartic acid | 17.49197 | 18.09886 | 1.0346953 | ns |
|  | phenylalanine | 0.851613 | 0.676995 | 0.7949562 | * |
| **Carbohydrates**  **(8)** | Cellobiose | 0.226552 | 0.241017 | 1.0638485 | ns |
|  | Xylose | 0.468207 | 0.525676 | 1.1227427 | ns |
|  | Fructose | 20.78008 | 12.59078 | 0.6059062 | *** |
|  | Mannose | 2.391477 | 2.10994 | 0.8822748 | ns |
|  | Galactinol | 6.276798 | 13.97621 | 2.2266465 | **** |
|  | Myo-inositol | 94.97775 | 168.9713 | 1.7790619 | **** |
|  | Sucrose | 623.9774 | 765.1692 | 1.2262771 | ns |
|  | Glucose | 203.0177 | 164.4232 | 0.8098959 | ns |
| **Fatty acids**  **(2)** | Lignoceric acid | 0.019337 | 0.014183 | 0.7334643 | ns |
|  | α-Linolenic acid | 1.023507 | 1.085415 | 1.0604862 | ns |
| **TCA intermediates**  **(4)** | Succinic acid | 7.641406 | 8.634707 | 1.1299893 | ns |
|  | Citric acid | 58.52785 | 88.8939 | 1.5188308 | **** |
|  | Fumaric acid | 0.086001 | 0.091698 | 1.0662434 | ns |
|  | Malic acid | 87.66116 | 130.8592 | 1.4927843 | **** |
| **Antioxidants**  **(3)** | α-tocopherol | 0.082851 | 0.16482 | 1.9893544 | *** |
|  | γ-tocopherol | 0.173117 | 0.221568 | 1.2798743 | * |
|  | Ascorbic acid | 0.808454 | 1.904183 | 2.3553387 | **** |
| **Others**  **(12)** | GABA | 1.228637 | 0.802649 | 0.6532841 | * |
|  | p-coumaric acid | 6.435078 | 5.587097 | 0.8682252 | ns |
|  | Glycerol | 2.488156 | 2.885107 | 1.1595362 | ns |
|  | Gluconic acid | 2.83886 | 2.714042 | 0.9560324 | ns |
|  | Ferulic acid | 0.409135 | 0.60419 | 1.4767497 | * |
|  | Glyceric acid | 5.384954 | 10.77265 | 2.0005092 | **** |
|  | Tartaric acid | 0.153954 | 0.190237 | 1.2356743 | ns |
|  | Threonic acid | 1.447445 | 1.891493 | 1.3067806 | ns |
|  | Phosphoric acid | 8.624856 | 6.335219 | 0.7345304 | ** |
|  | Phytol | 0.22284 | 0.34615 | 1.5533567 | *** |
|  | Urea | 0.22211 | 0.612894 | 2.7594165 | ** |
|  | Coutraric acid | 3.585422 | 4.25343 | 1.1863122 | ns |

^a^ns = not significant.

^b^*, **, ***, and **** denotes statistical significance at the p < 0.05, 0.01, 0.001, and 0.0001 levels, respectively, as per Student's *t*-tes

**Supplementary Table 2.** Loading values for 46 metabolites in spinach leaves as separated by two PCA components

| **Metabolites** | **PC1 (37.4 %)** | **Metabolites** | **PC2 (18.2 %)** |
| --- | --- | --- | --- |
|  | **Loading values** |  | **Loading values** |
| Urea | 0.424740445 | Urea | 0.343027278 |
| Leucine | 0.326670806 | 5-oxoproline | 0.242793877 |
| Proline | 0.298059641 | α-tocopherol | 0.237116353 |
| α-tocopherol | 0.264255115 | Xylose | 0.236380624 |
| Methionine | 0.259850001 | Coutaric acid | 0.196370915 |
| Galactinol | 0.247665957 | Ferulic acid | 0.171065458 |
| Ascorbic acid | 0.240624619 | Glucose | 0.168953071 |
| Glyceric acid | 0.199585851 | Tartaric acid | 0.138737141 |
| Glycine | 0.198521901 | Phenylalanine | 0.138210121 |
| Phytol | 0.174829047 | Fructose | 0.128048775 |
| Myo-inositol | 0.173717913 | GABA | 0.110315225 |
| Cysteine | 0.147217617 | Phosphoric acid | 0.100964800 |
| Ferulic acid | 0.127791038 | α-linolenic acid | 0.094290753 |
| Threonine | 0.122881210 | Phytol | 0.085488159 |
| Malic acid | 0.115069704 | β-alanine | 0.071050828 |
| Coutaric acid | 0.110884777 | Isoleucine | 0.067868768 |
| Citric acid | 0.110053031 | Valine | 0.053299247 |
| Glutamine | 0.106313321 | Coumaric acid | 0.043453650 |
| Glutamic acid | 0.099337917 | Glycerol | 0.029939064 |
| Isoleucine | 0.096002884 | Glyceric acid | 0.004259398 |
| γ-tocopherol | 0.084776527 | Succinic acid | 0.002063015 |
| 5-oxoproline | 0.081245217 | Aspartic acid | -0.002618612 |
| Xylose | 0.074080294 | Lignoceric acid | -0.006468998 |
| Glycerol | 0.072209379 | Methionine | -0.009586754 |
| Tartaric acid | 0.070417794 | Malic acid | -0.011043896 |
| Threonic acid | 0.068581664 | Tyrosine | -0.011501021 |
| Sucrose | 0.058958705 | Citric acid | -0.012915525 |
| Valine | 0.056613685 | Myo-inositol | -0.034091772 |
| β-alanine | 0.049560627 | Galactinol | -0.034843029 |
| Succinic acid | 0.036168074 | Serine | -0.036668078 |
| Serine | 0.032801228 | Glycine | -0.045984952 |
| α-linolenic acid | 0.032430298 | γ-tocopherol | -0.053902262 |
| Tyrosine | 0.025741058 | Mannose | -0.085402033 |
| Aspartic acid | 0.007932764 | Glutamic acid | -0.100570183 |
| Cellobiose | -0.004886460 | Proline | -0.103886251 |
| Gluconic acid | -0.011870800 | Ascorbic acid | -0.104643241 |
| Fumaric acid | -0.022669038 | Gluconic acid | -0.106779455 |
| Coumaric acid | -0.036094160 | Threonine | -0.152738902 |
| Glucose | -0.040038972 | Asparagine | -0.153872396 |
| Lignoceric acid | -0.048882012 | Threonic acid | -0.155846216 |
| Mannose | -0.053079655 | Glutamine | -0.158245061 |
| Phenylalanine | -0.055071995 | Fumaric acid | -0.182104311 |
| Phosphoric acid | -0.069639111 | Cysteine | -0.194649534 |
| Fructose | -0.108119987 | Sucrose | -0.223432340 |
| Asparagine | -0.123123022 | Cellobiose | -0.291352026 |
| GABA | -0.123893210 | Leucine | -0.379807424 |
